# Segregation of pathways leading to pexophagy

**DOI:** 10.1101/2022.11.08.515582

**Authors:** Francesco G. Barone, Sylvie Urbé, Michael J. Clague

## Abstract

Peroxisomes are organelles with key roles in metabolism including long-chain fatty acid production. Their metabolic functions overlap and interconnect with mitochondria, with which they share an overlapping but distinct proteome. Both organelles are degraded by selective autophagy processes termed pexophagy and mitophagy. Whilst mitophagy has received intense attention, the pathways linked to pexophagy and associated tools are less well developed. We have identified the neddylation inhibitor, MLN4924, as a potent activator of pexophagy and show that this is mediated by the HIF1α-dependent upregulation of BNIP3L/NIX, a known adaptor for mitophagy. We show that this pathway is distinct from pexophagy induced by the USP30 deubiquitylase inhibitor, CMPD-39, for which we identify the adaptor NBR1 as a central player. Our work suggests a level of complexity to the regulation of peroxisome turnover that includes the capacity to co-ordinate with mitophagy, via NIX, which acts as a rheostat for both processes.

## Introduction

Selective autophagy requires that an organelle destined for elimination is marked and then linked to the phagophore membrane. This normally occurs via a LC3-interacting region (LIR) that links to lipid modified LC3 (1,2). In many cases these selective autophagy adaptors are recruited to damaged organelles by binding to ubiquitin, which accumulates on their surface. These include adaptors from the Sequestosome-1-like receptor family that include p62, NBR1, NDP52, TAX1BP1 and OPTN. All of these proteins have both a LIR and a ubiquitin binding domain. Nevertheless, they are differentially associated with various forms of selective autophagy and the underlying reasons are not completely understood (3). For example, a comparison of these factors in the regulation of ubiquitin-dependent mitophagy suggested that only NDP52 and OPTN play critical roles (4). In contrast, ubiquitin-dependent pexophagy relies upon NBR1 (5,6). There is also diversity of mechanism in pathways converging on organelle turnover. For example, mitophagy is induced by mitochondrial depolarisation, activating the PINK1-PRKN pathway which leads to coating the outer mitochondrial surface with ubiquitin and is suppressed by the deubiquitylase (DUB), USP30 (7,8). However, the vast majority of mitophagy in an organism is PINK1-PRKN independent (9,10). An alternative ubiquitin-independent pathway exists, that employs trans-membrane domain containing proteins, BNIP3 and BNIP3L/NIX, which insert directly into the polarised mitochondrial membrane and link to the phagophore LC3 by LIR domains (11). NIX plays a critical role in the removal of mitochondria during reticulocyte development and neuronal differentiation and is up-regulated by hypoxia (12–16).

Inherited mutations in peroxisomal genes can lead to debilitating peroxisomal disorders. Many of these have been linked to peroxisome biogenesis, but it is now apparent that the other arm of peroxisome homeostasis, namely pexophagy, is also critical (17,18). For example, the AAA-ATPase comprised of PEX1, PEX6 and PEX26 is a pexophagy suppressor by virtue of removing the ubiquitylated peroxisomal matrix protein import receptor, Ub-PEX5 from the peroxisomal membrane (19). This shift into the spotlight has highlighted an unmet need for tools to manipulate pexophagy in a controlled manner. Here we started out surveying a number of candidate compounds and discovered that the neddylation inhibitor MLN4924 and the USP30 inhibitor, compound 39 (CMPD-39) are both potent inducers of pexophagy (20,21). In a parallel to mitophagy, we show that the MLN4924 effect reflects activation of HIF1α and upregulation of NIX. In contrast, CMPD-39 promotes a pexophagy pathway that uses the ubiquitin recognising adaptor NBR1, consistent with the proposed mode of action on a DUB previously linked to pexophagy (22,23). Our results highlight the increasing awareness of the complexity of pexophagy pathways akin to mitophagy and suggest that, in the case of hypoxia/MLN4924 treatment, both organelles can be turned over in a co-ordinated manner.

## Results

### Survey of pexophagy inducing agents

We generated a retinal pigment epithelial (hTERT-RPE1) cell line, expressing a peroxisomal matrix targeting signal joined with a mKeima fluorophore (Keima-SKL). This fluorophore reports on lysosomal delivery of peroxisomes. The lower pH of the lysosome leads to a change in the excitation spectrum properties of the reporter and the resultant pexolysosomes are represented in red pseudocolour (Figure 1A) (22,24). We conducted a survey of various chemicals, drawn from the literature, for their influence on pexophagy. These included agents linked to peroxisome turnover (4-phenylbutyric acid - 4-PBA, Clofibrate), others previously linked to mitophagy (Deferiprone - DFP, MLN4924) and the USP30 inhibitor CMPD-39, which we have previously shown to promote both pathways as well as a general oxidative stress (21,25–27).

**Figure 1.**
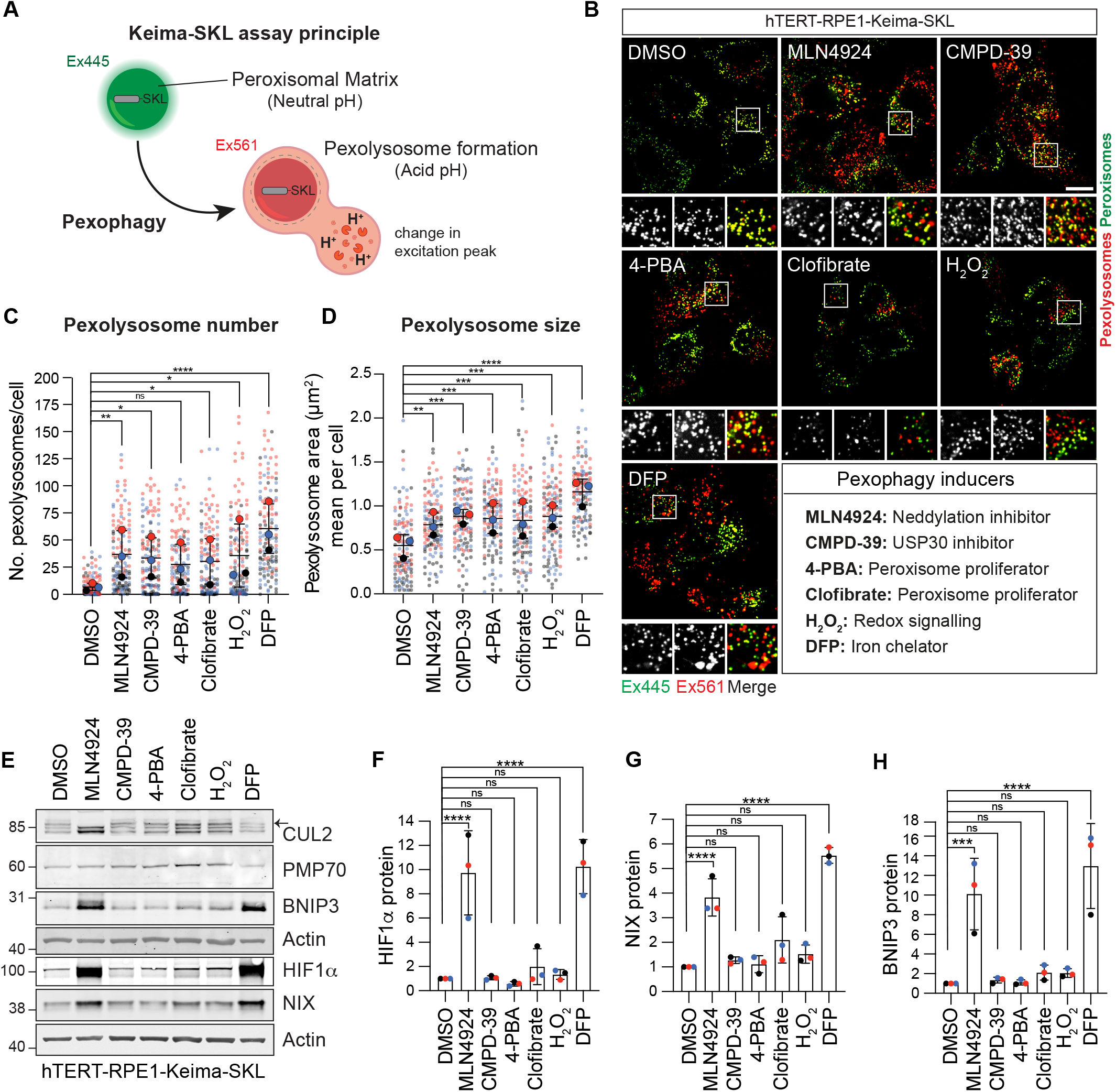
A chemical screen for pexophagy inducers. **(A)** Schematic for the Keima-SKL pexophagy reporter system. The Keima fluorescent reporter is targeted to the peroxisomal matrix via the peroxisomal targeting signal 1 (tripeptide SKL). Upon delivery to the acidic environment of lysosomes, the excitation spectrum of Keima is red shifted. Fluorescence emission from these two excitation wavelengths (445 and 561 nm) is pseudocoloured green and red respectively **(B)** Representative images of hTERT-RPE1 cells stably expressing the Keima-SKL pexophagy reporter. CMPD-39 (1 μM) was administered for 96 h before imaging. For all other conditions, cells were treated for 24 hours with MLN4924 (1 μM), 4-PBA (4-PBA, 1 mM), Clofibrate (20 μM), hydrogen peroxide (H_2_O_2_, 100 μM) and Deferiprone (DFP, 1 mM). Scale bar 20 μm. **(CD)** Graphs illustrate the number and mean area of pexolysosomes per cell. Quantification of the data from three independent colour coded experiments is shown. Mean and standard deviation (SD) are indicated; >40 cells were quantified per condition in each replicate experiment. One way ANOVA and Bonferroni’s multiple comparison test. *P<0.05, **P<0.01, ***P<0.001, ****P<0.0001 **(E)** Representative Western blot of hTERT-RPE1-Keima-SKL treated as in (B), probed for Cullin-2 (CUL2), PMP70, BNIP3, HIF1α, NIX and Actin. Arrow indicates the neddylated form of Cullin-2. **(F-G-H)** Quantitation of data shown in (E) indicating the mean and standard deviation for three independent colour coded experiments.

All agents led to increases in pexophagy indices of similar magnitudes (Figure 1B-D, Supplementary Figure 1A-C). Both DFP and MLN4924 generate an increase in HIF1α, which in turn drives expression of the mitophagy adaptors, BNIP3 and NIX (Figure 1E-H). DFP inhibits the prolyl hydroxylase enzyme that renders HIF1α a substrate for the Cullin RING ligase VHL, whilst MLN4924 inhibits the neddylation-dependent activation of all Cullins (Figure 1E). We have subsequently focused our efforts on characterising CMPD-39 and MLN4924 because of their high selectivity for their protein targets.

### Elevated NIX levels promote pexophagy

We next showed that MLN4924-induced pexophagy requires the canonical autophagy machinery as the number of pexolysosomes is sensitive to depletion of the key autophagy orchestrator ATG7 (Figure 2A-D). It is insensitive to depletion of the peroxisomal membrane protein ACBD5, whose yeast counterpart, Atg37, has been strongly linked to pexophagy (28). However, the combined depletion of BNIP3 and NIX restores pexophagy to baseline levels suggesting that these proteins fully account for the observed effect of MLN4924 (Figure 2A-D). Individual depletions of BNIP3 and NIX indicates that NIX is the principal inducer of pexophagy (Figure 2E-H).

**Figure 2:**
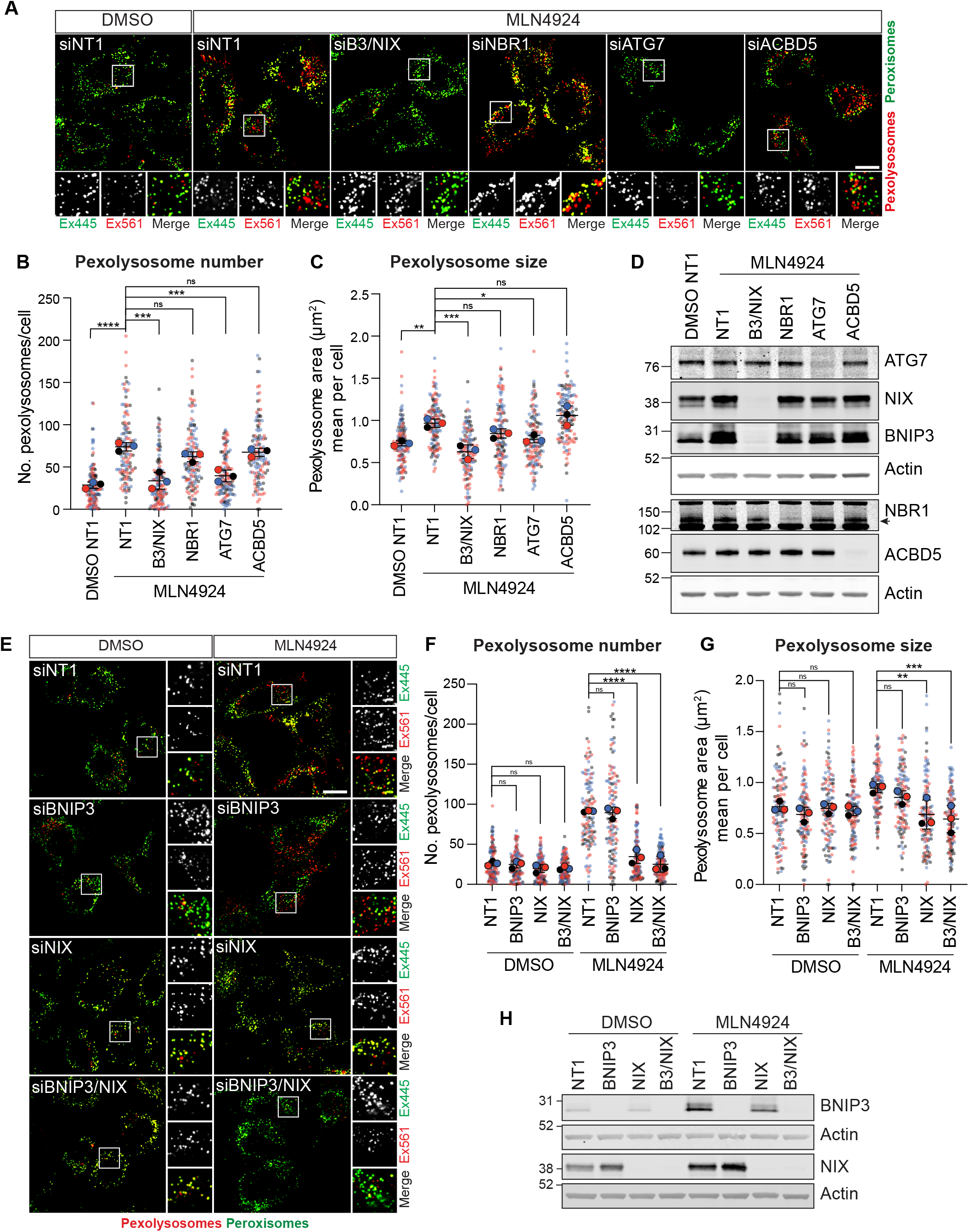
MLN4924-induced pexophagy requires NIX. **(A)** Representative confocal images of hTERT-RPE1-Keima-SKL cells treated with DMSO and MLN4924 (1 μM) for 24 h before imaging. Cells were transfected with non-targeting siRNA (NT1) or siRNA targeting BNIP3, NIX, ATG7, ACBD5 and NBR1. Scale bar 20 μm. **(B-C)** Graphs show the number and mean area of pexolysosomes. Quantification of the data from three colour coded independent experiments is shown. Mean and SD are indicated; >40 cells were quantified per condition in each experiment. One-way ANOVA with Bonferroni’s multiple comparisons test. *P < 0.05. **P < 0.01. ***P < 0.001. ****P < 0.0001. **(D)** Representative Western blot of hTERT-RPE1-Keima-SKL cells treated as in (A) and probed as indicated. **(E)** Representative confocal images of hTERT-RPE1-Keima-SKL cells stably expressing the Keima-SKL pexophagy reporter. Cells were treated with DMSO and MLN4924 (1 μM), for 24 h before imaging. Cells were transfected with non-targeting (NT1) siRNA or siRNA targeting BNIP3 and NIX. Scale bar 20 μm. **(F-G)** Graph shows the number and mean area of pexolysosomes per cell. Quantification of the data from three colour-coded independent experiments is shown. Mean and standard deviation are indicated; >40 cells were quantified per condition in each repeat experiment. One-way ANOVA with Bonferroni’s multiple comparisons test. **P < 0.01. ***P < 0.001. ****P < 0.0001 **(H)** Representative Western blot of protein samples from cells treated as in (E) and probed as indicated.

### Direct association of NIX with peroxisomes

Whilst NIX is known to directly insert into the mitochondrial membrane, its association with peroxisomes has not been previously established. We wished to check if we could observe NIX on peroxisomes and secondly whether this reflects a direct association or transfer from mitochondria. We employed a strategy we have previously used to demonstrate insertion of USP30 directly into the peroxisomal membrane (22). We elected to effectively remove mitochondria from hTERT-RPE1 cells, stably expressing high levels of YFP-Parkin, by eliciting mitophagy following mitochondrial depolarisation with antimycin and oligomycin (A/O, Figure 3A). Subsequently, cells were transfected with DsRed-NIX and examined by fluorescence microscopy for colocalisation with the peroxisomal marker PMP70 (Figure 3B,C) or harvested for subcellular fractionation (Figure 3D). We find clear evidence for direct association of NIX with peroxisomal membranes under these conditions, suggesting that the effect of NIX on pexophagy reflects this association rather than some indirect consequence of mitophagy.

**Figure 3.**
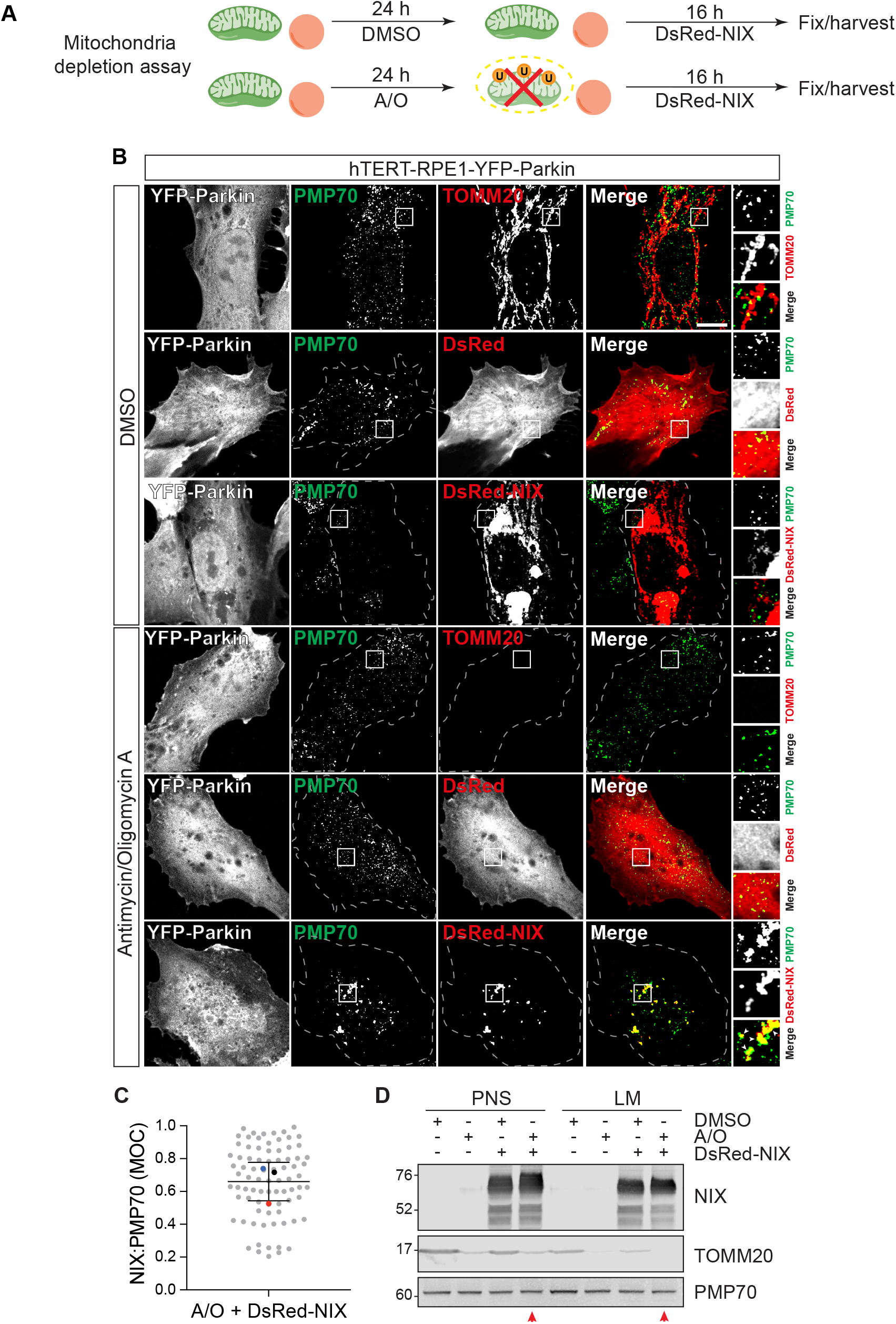
Exogenously expressed NIX colocalises with peroxisomes independently of mitochondria. **(A)** hTERT-RPE1-YFP-Parkin cells were first treated for 24 h with oligomycin A and antimycin A (AO,1 μM each) or DMSO, then transiently transfected with DsRed-NIX or DsRed alone for 16 h prior to fixation or harvesting. **(B)** Representative images of cells, treated as described in (A), immunostained for endogenous PMP70 (AlexaFluor-405, green). A set of mock transfected cells treated in parallel were co-stained for TOMM20 (AlexaFluor-594, red). Scale bars 10 μm. **(C)** Colocalisation of the DsRed-NIX signal with peroxisomes (PMP70) in hTERT-RPE1-YFP-Parkin cells treated with A/O. The Manders overlap coefficient (MOC, M1) was calculated for >25 cells per experiment. The mean and standard deviation of three independent experiments is shown. **(D)** Representative Western blot showing subcellular fractions of hTERT-RPE1-YFP-Parkin cells treated as shown in (A). Post-nuclear supernatant (PNS) and light membranes (LM) are shown; representative of two independent experiments, red arrow indicates the mitochondria-depleted, DsRed-NIX transfected PNS and LM samples.

### CMPD-39 induced pexophagy requires NBR1

Whilst NIX expression is under the control of ubiquitin E3-ligase activity, the actual pexophagy event does not require organelle coating with ubiquitin, owing to its direct insertion into membranes. Moreover, CMPD-39 does not increase NIX levels (Figure 1E,G). We presumed that USP30 inhibition by CMPD-39 must be promoting autophagy via an alternative ubiquitin-dependent pathway. We thus chose to test a role for the established pexophagy adaptor NBR1 in this system, which acts as a bridge between ubiquitin and LC3. The increased pexophagy following CMPD-39 is insensitive to BNIP3/NIX or ACBD5 depletion but returns to baseline on depletion of ATG7 or NBR1 (Figure 4A-D).

**Figure 4.**
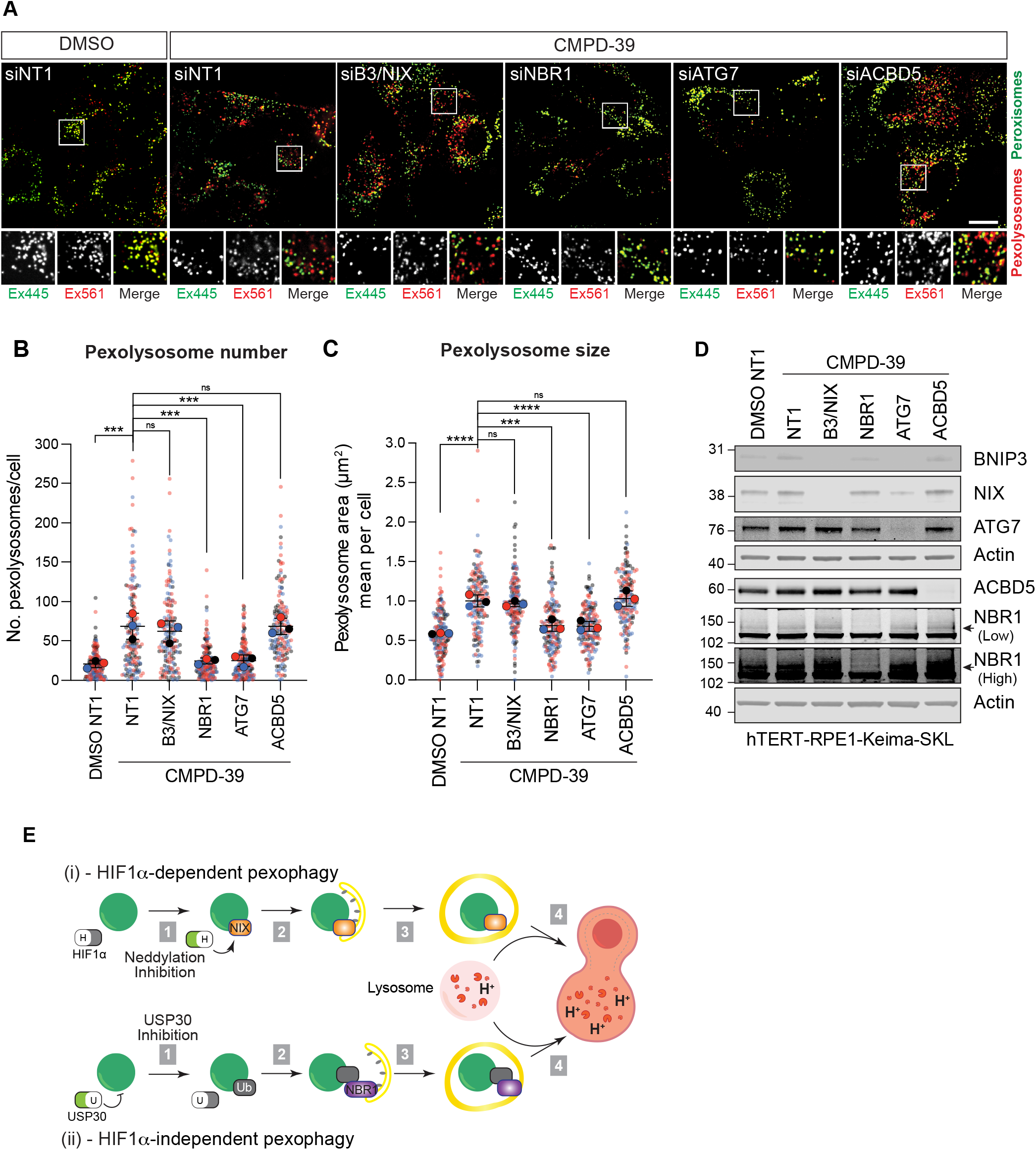
NBR1 mediates CMPD-39-induced pexophagy. **(A)** Representative confocal images of hTERT-RPE1-Keima-SKL cells treated with DMSO and CMPD-39 (1 μM), for 96 h before imaging. Cells were transfected with non-targeting (NT1) or siRNA or siRNA targeting BNIP3, NIX, ATG7, ACBD5 and NBR1. Scale bar 20 μm. Quantification of data shown in A. **(B,C)** Graphs show the number and mean area of pexolysosomes per cell from three independent colour coded experiments. Mean and standard deviation are indicated; >50 cells were quantified per condition in each experiment. One-way ANOVA with Bonferroni’s multiple comparisons test. ***P < 0.001. ****P < 0.0001. **(D)** Representative Western blot of protein samples from cells treated as in (A) and probed as indicated. Low and high represent two different exposures of the same blot. **(E)** HIF1α-dependent and -independent pexophagy pathways. (i) HIF1α-dependent pexophagy pathway. Upon administration of the neddylation inhibitor, MLN4924, the transcription factor HIF1α is induced (top). This leads to the upregulation of NIX, which directly associates with peroxisomes and acts as a pexophagy adaptor. (ii) HIF1α-independent pexophagy pathway: the DUB USP30 suppresses pexophagy by removing ubiquitin attached to peroxisomal substrates. Accumulation of ubiquitin on peroxisomes, following USP30 inhibition, leads to the recruitment of the NBR1 pexophagy adaptor.

## Discussion

Here we have used MLN4924 to induce NIX via the HIF1α pathway (Figure 4E). We show that NIX is targeted to peroxisomes and promotes pexophagy. This provides an additional function for NIX beyond its established role in governing mitophagy in various contexts (12,14,15,29,30). Whilst this manuscript was in preparation, Ganley and colleagues reported similar findings, using the iron chelating drug DFP, to induce HIF1α and hence NIX (31). In our hands DFP has a stronger pexophagy-promoting effect than MLN4924 despite the latter’s complete inhibition of neddylation. As the CRL2^VHL^ complex is the principal regulator of HIF1α stability and requires neddylation for activity, we suggest that there may be additional ill-defined pathways induced by DFP (32). Both drugs provide a complementary approach to inducing pexophagy and can be usefully cross-referenced with each other in future studies. MLN4924 (otherwise known as Pevonedistat) is well tolerated and has featured in more than thirty oncology related clinical trials (33). In this disease context, up-regulation of the HIF1α pathway is not desirable and our work highlights that consideration should be given to combination therapy together with HIF pathway inhibitors.

It has been estimated that 2/3 of the respective proteomes of mitochondria and peroxisomes overlap (34). There are close contacts between the two organelles and some mitochondrial material can be delivered to peroxisomes via vesicular transport (35). We have previously shown that USP30 can be directly targeted to peroxisomes in cells lacking mitochondria (22). We now show that the same holds true for NIX. Thus the effect of NIX on pexophagy does not require its traversal through mitochondria.

We have also previously shown CMPD-39 induces pexophagy (21). Here, we have been able to compare the magnitude of this effect with other agents. In our survey of pexophagy inducing chemicals, CMPD-39 induced pexophagy to a similar degree as MLN4924. However, CMPD-39 induced pexophagy is entirely dependent on NBR1 and this distinguishes it from the MLN4924-NIX pathway (Figure 4E). The ubiquitin-binding pexophagy adaptor, NBR1, has previously been shown to be necessary and sufficient for basal pexophagy (5,36). The simplest model predicts that peroxisome localised USP30 suppresses critical ubiquitin signals that otherwise lead to pexophagy (22,23). This concords with the proposed role of USP30 in mitophagy, although the downstream adaptor may vary (22,30,37–43). Riccio and colleagues have suggested that USP30 suppresses peroxisome ubiquitylation attributed to the peroxisomal E3 ligase PEX2 (23). More recently Zheng and colleagues have shown that the E3 ligase MARCH5 is shared between mitochondria and peroxisomes and can also promote pexophagy (44). At mitochondria, MARCH5 and USP30 have been shown to reciprocally regulate the ubiquitin status of the mitochondrial import (TOMM) complex (41). We have proposed that this sets a trigger threshold for unleashing the PINK1-PRKN cascade (22,42,45). We speculate that a MARCH5/USP30 counterpoise system may be preserved at the peroxisome and is disrupted by CMPD-39. Both pexophagy pathways, whose induction we have chemically segregated here, have key factors in common with established mitophagy pathways (Figure 4E). Given the metabolic interplay between these two organelles, one can imagine advantages in regulating their abundance in a co-ordinated manner.

## Materials and Methods

### Cell culture, transfection and RNA interference

hTERT-RPE1-Keima-SKL and hTERT-RPE1-YFP-Parkin cells were routinely cultured in Dulbecco’s Modified Eagle’s medium DMEM/F12 (Gibco, 31331028) supplemented with 10% FBS (Gibco, 10270106), 1% non-essential amino acids (Gibco, 111505035) and 1% penicillin/streptomycin at 37°C and 5% CO_2_. Cells were routinely checked for mycoplasma. For RNA interference experiments, cells were treated with 40 nM of non-targeting (NT1) or target specific siRNA oligonucletotides (Dharmacon On-target plus smart pool), using Lipofectamine RNAi-MAX (Invitrogen, 13778030) according to manufacturer’s instructions. For plasmid transfections, Lipofectamine 3000 (Invitrogen, L3000001) was used according to the manufacturers instructions.

### Generation of the pexophagy reporter hTERT-RPE1-Keima-SKL cell line

hTERT-RPE1-Cas9i-PuroS cells (46), were transfected with pCDNA3.1-mKeima-SKL-BlastR (derived from pCDNA3.1-mKeima-SKL-Neo (22)), using Lipofectamine 2000 (Invitrogen, 11668019). Transfected cells were selected with 10 μg/ml Blasticidin S HCl for 7 days and mKeima positive cells were isolated by FACS. A clonally isolated cell line expressing suitable levels of the mKeima-SKL reporter was selected and is referred to as hTERT-RPE1-Keima-SKL for all experiments in this paper.

### siRNA and plasmids

The following ON-Target Plus Smart Pool siRNAs were obtained from Dharmacon: BNIP3 (5’ -UCGCAGACACCACAAGAUA-3’, 5’ -GAACUGCACUUCAGCAAUA-3’, 5’ -GGAAAGAAGUUGAAAGCAU-3’, 5’ -ACACGAGCGUCAUGAAGAA-3’), BNIP3L/NIX (5’ -GACCAUAGCUCUCAGUCAG-3’, 5’ -CAACAACAACUGCGAGGAA-3’, 5’ -GAAGGAAGUCGAGGCUUUG-3’, 5’ -GAGAAUUGUUUCAGAGUUA-3’), ATG7 (5’ -CCAACACACUCGAGUCUUU-3’, 5’ -GAUCUAAAUCUCAAACUGA-3’, 5’ -GCCCACAGAUGGAGUAGCA-3’, 5’ -GCCAGAGGAUUCAACAUGA-3’. ACBD5 (5’ -CUAAAGGGAUCUACUACUA-3’, 5’ -CCAAAACCGUUAAUGGUAA-3’, 5’ -CAGCAUUUGACAAGCGAUU-3’, 5’ -GGAUGCAACACUUGAGCGA-3’. NBR1 (5’ -GAGAACAAGUGGUUAACGA-3’, 5’ -CCACAUGACAGUCCUUUAA-3’, 5’ -GAACGUAUACUUCCCAUUG-3’, 5’ -AGAAGCCACUUGCACAUUA-3’. DsRed-BNIP3L/NIX plasmid was obtained from Addgene (100763). DsRed-N1 plasmid was a gift from Francis Barr (University of Oxford).

### Antibodies and reagents

Antibodies and other reagents used were as follows: anti-BNIP3L (Cell Signalling, #12396, 1:1000 WB), anti-BNIP3 (Abcam, ab109362, 1:1000 WB), anti-TOM20 (ProteinTech, 11802-1-AP, 1:1000 WB), anti-TOM20 (Sigma, HPA011562, 1:500 IF), anti-PMP70 (Sigma, SAB4200181, 1:1000 WB, 1:250 IF), mouse anti-actin (ProteinTech, 66009-1-Ig, 1:10,000), anti-Cullin2 (Bethyl, A302-476A, 1:2000 WB), anti-HIF1α (NB100-134, 1:1000, Novus Bio techne), anti-ATG7 (Cell Signalling, 2631, 1:1000 WB), anti-NBR1 (Cell Signalling, 9891S, 1:1000 WB), MLN4924 (Chemgood, C-1231), Deferiprone (Sigma, 379409), 4-PBA (Sigma, 21005), Clofibrate (Sigma, C6643), Hydrogen peroxide (Sigma, H1009), oligomycin A (Sigma, 75351), antimycin A (Sigma, A8674).

### Preparation of cell lysates and Western blot analysis

Cultured cells were lysed with RIPA (150 mM Sodium chloride (Fisher Scientific, S316060), 1% Sodium deoxycholate, 0.1% SDS, 1% Triton X-100) lysis buffer and routinely supplemented with mammalian protease inhibitor cocktail (Sigma, P8340) and Phostop (Roche, 04906837001). Proteins were resolved using SDS–PAGE (Invitrogen NuPage gel 4–12%, NP0321/NP0335), transferred to nitrocellulose membrane (Amersham, 10600002), blocked in 5% milk (Marvel) or 5% BSA (First Link, 41-10-410) in TBS (20mM Tris-Cl pH 7.6, 150mM NaCl) supplemented with Tween-20 (Fisher Scientific, 10485733), and probed with primary antibodies overnight. Visualisation and quantification of Western blots were performed using IRdye 800CW and 680LT coupled secondary antibodies and an Odyssey infrared scanner (LI-COR Biosciences, Lincoln, NE).

### Mitochondria depletion assay

As previously described (22), hTERT-RPE1 cells stably overexpressing YFP-Parkin (hTERT-RPE1-YFP-Parkin) were treated for 24 h with Antimycin A (1 μM) and Oligomycin A (1 μM), washed with PBS and cultured in normal media prior to transfection with a plasmid expressing DsRED alone or DsRed-BNIP3L/NIX (22). After 16 hours, cells were fixed for immunofluorescence microscopy or harvested for subcellular fractionation.

### Subcellular fractionation

hTERT-RPE1-YFP-Parkin cells were washed twice with ice-cold PBS, followed by centrifugation at 1000g for 2 min at 4°C. Cell pellets were resuspended in HIM buffer (200 mM Mannitol, 70 mM Sucrose, 1mM EGTA and 10 mM Hepes-NaOH, pH 7.4) and centrifuged again at 1000g for 5 min at 4°C. The cell pellet was resuspended in HIM buffer, supplemented with 50 mM 2-Chloroacetamide, mammalian protease inhibitor cocktail (Sigma, P8340) and Phostop (Roche, 04906837001), before cells were mechanically disrupted by shearing with a 23 gauge needle. The cell homogenate was then centrifuged at 600 g for 10 min to obtain the post-nuclear supernatant (PNS). Heavy membranes enriched in mitochondria were removed by centrifugation at 7000 g for 15 min and the supernatant was centrifuged at 100,000 g for 30 min to generate the light membrane pellet fraction (LM). Equal amount of protein from both PNS and LM (10 μg/lane) was resolved by SDS-PAGE (Invitrogen NuPage gel 4–12%).

### Immunofluorescence and colocalization analysis

hTERT-RPE1-YFP-Parkin cells were fixed using 4% paraformaldehyde in PBS, permeabilised with 0.2% Triton X-100 in PBS, stained with AlexaFluor-405 or −594 coupled secondary antibodies and imaged using a Zeiss LSM900 with Airyscan (63× NA 1.4 oil, acquisition software Zen Blue). The images were processed using Adobe Photoshop 2022 and Fiji v2.9.0 software. For colocalisation analysis, single confocal z-planes were analysed with the JaCoP plugin in Fiji v2.9.0 to derive the Manders overlap coefficient (MOC) M1. Quantification of colocalisation was performed from three independent experiments analysing >25 cells per experiment.

### Live cell imaging of pexophagy

For live cell imaging, hTERT-RPE1-Keima-SKL cells were seeded onto an IBIDI μ-Dish (2×10^5^) (IBIDI, 81156) two days before image acquisition using a 3i Marianas spinning disk confocal microscope (63x oil objective, NA 1.4, Photometrics Evolve EMCCD camera, Slide Book 3i v3.0). Live cells were imaged sequentially (Ex445/Em600 then Ex561/Em600). Images were processed using Adobe Photoshop 2022 and Fiji v2.9.0 softwares. Analysis of pexophagy levels in hTERT-RPE1-Keima-SKL was performed using the semi-automated ‘mito-QC Counter’ plugin implemented in Fiji v2.9.0 software as previously described (47). The analysis of pexophagy was performed for three independent experiments analysing >40 cells per condition in each experiment.

### Statistical analysis

P-values are indicated as *P<0.05, **P<0.01, ***P<0.001, ****P<0.0001 and derived by one-way ANOVA and Bonferroni’s multiple comparisons post hoc test. All statistical analysis was conducted using GraphPad Prism 9.

## Acknowledgements

hTERT-RPE1-YFP-Parkin cells were a kind gift of Jon Lane (University of Bristol). FGB was funded by a Wellcome Trust PhD studentship, 102172/B/13/Z. MJC is a Royal Society Industry Fellow, INF\R2\212031. Additional support was provided by Bristol-Myers Squibb.

## Figure Legends

**Supplementary Figure 1.**
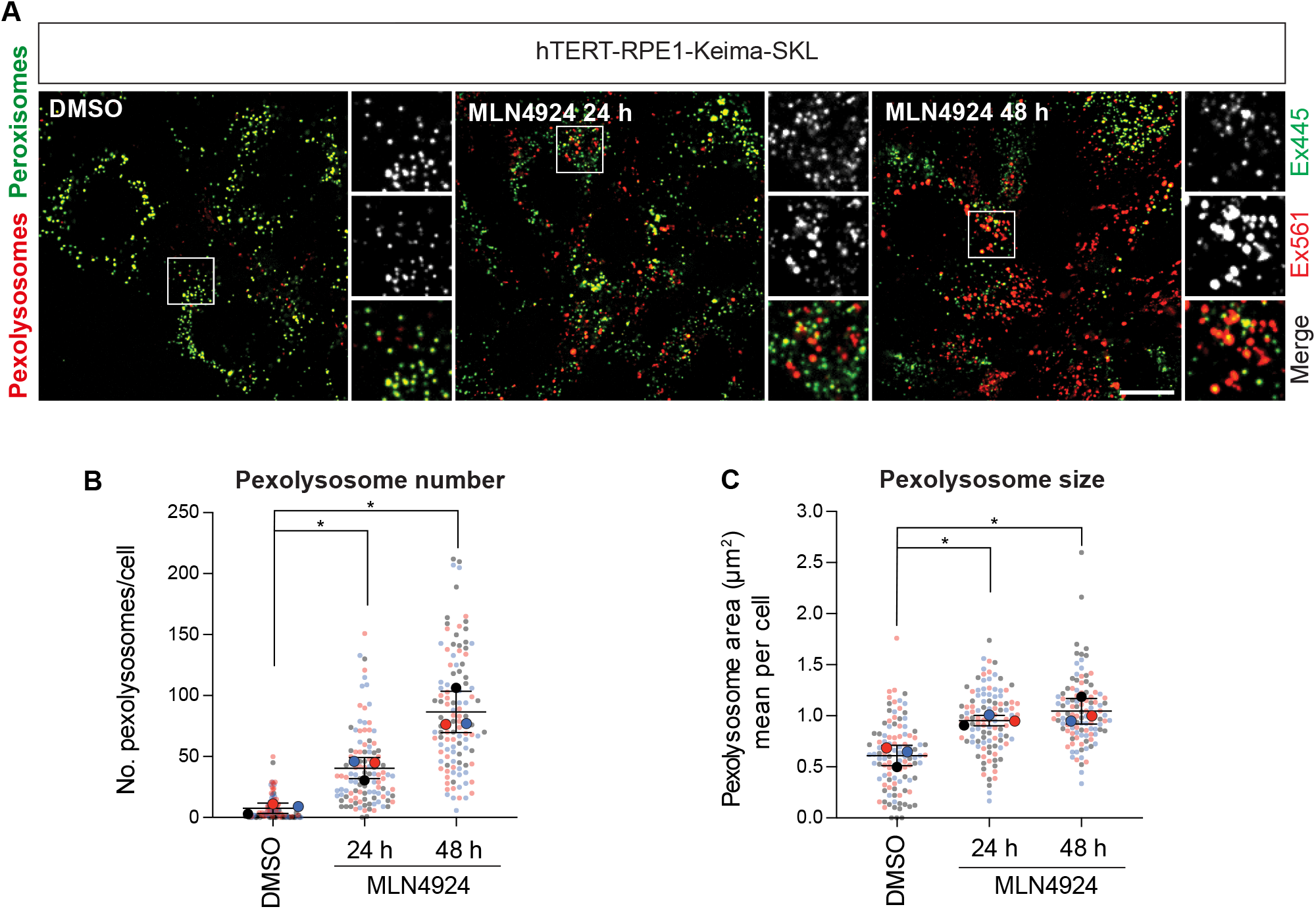
Neddylation inhibition induces pexophagy. **(A)** Representative images of hTERT-RPE1-Keima-SKL cells. MLN4924 (1 μM) was administered for 24 and 48 h before imaging. Scale bar 20 μm. (B-C) Graphs show the number and mean area of pexolysosomes. Quantification of the data from three independent colour coded experiments is shown. Mean and standard deviation are indicated; >30 cells were quantified per condition in each experiment. One-way ANOVA with Bonferroni’s multiple comparisons test. *P < 0.05.

## References

1. Johansen, T., and Lamark, T. (2020) Selective Autophagy: ATG8 Family Proteins, LIR Motifs and Cargo Receptors. J Mol Biol 432, 80–103

2. Lamark, T., and Johansen, T. (2021) Mechanisms of Selective Autophagy. Annu Rev Cell Dev Biol 37, 143–169

3. Gubas, A., and Dikic, I. (2022) A guide to the regulation of selective autophagy receptors. FEBS J 289, 75–89

4. Lazarou, M., Sliter, D. A., Kane, L. A., Sarraf, S. A., Wang, C., Burman, J. L., Sideris, D. P., Fogel, A. I., and Youle, R. J. (2015) The ubiquitin kinase PINK1 recruits autophagy receptors to induce mitophagy. Nature 524, 309–314

5. Deosaran, E., Larsen, K. B., Hua, R., Sargent, G., Wang, Y., Kim, S., Lamark, T., Jauregui, M., Law, K., Lippincott-Schwartz, J., Brech, A., Johansen, T., and Kim, P. K. (2013) NBR1 acts as an autophagy receptor for peroxisomes. J Cell Sci 126, 939–952

6. Rasmussen, N. L., Kournoutis, A., Lamark, T., and Johansen, T. (2022) NBR1: The archetypal selective autophagy receptor. J Cell Biol 221

7. Bingol, B., and Sheng, M. (2016) Mechanisms of mitophagy: PINK1, Parkin, USP30 and beyond. Free Radic Biol Med 100, 210–222

8. Pickles, S., Vigie, P., and Youle, R. J. (2018) Mitophagy and Quality Control Mechanisms in Mitochondrial Maintenance. Curr Biol 28, R170–R185

9. Lee, J. J., Sanchez-Martinez, A., Zarate, A. M., Beninca, C., Mayor, U., Clague, M. J., and Whitworth, A. J. (2018) Basal mitophagy is widespread in Drosophila but minimally affected by loss of Pink1 or parkin. J Cell Biol 217, 1613–1622

10. McWilliams, T. G., Prescott, A. R., Montava-Garriga, L., Ball, G., Singh, F., Barini, E., Muqit, M. M. K., Brooks, S. P., and Ganley, I. G. (2018) Basal Mitophagy Occurs Independently of PINK1 in Mouse Tissues of High Metabolic Demand. Cell Metab 27, 439–449 e435

11. Ney, P. A. (2015) Mitochondrial autophagy: Origins, significance, and role of BNIP3 and NIX. Biochim Biophys Acta 1853, 2775–2783

12. Schweers, R. L., Zhang, J., Randall, M. S., Loyd, M. R., Li, W., Dorsey, F. C., Kundu, M., Opferman, J. T., Cleveland, J. L., Miller, J. L., and Ney, P. A. (2007) NIX is required for programmed mitochondrial clearance during reticulocyte maturation. Proc Natl Acad Sci U S A 104, 19500–19505

13. Mazure, N. M., and Pouyssegur, J. (2010) Hypoxia-induced autophagy: cell death or cell survival? Curr Opin Cell Biol 22, 177–180

14. Zhang, J., and Ney, P. A. (2010) Reticulocyte mitophagy: monitoring mitochondrial clearance in a mammalian model. Autophagy 6, 405–408

15. Esteban-Martinez, L., Sierra-Filardi, E., McGreal, R. S., Salazar-Roa, M., Marino, G., Seco, E., Durand, S., Enot, D., Grana, O., Malumbres, M., Cvekl, A., Cuervo, A. M., Kroemer, G., and Boya, P. (2017) Programmed mitophagy is essential for the glycolytic switch during cell differentiation. EMBO J 36, 1688–1706

16. Ordureau, A., Kraus, F., Zhang, J., An, H., Park, S., Ahfeldt, T., Paulo, J. A., and Harper, J. W. (2021) Temporal proteomics during neurogenesis reveals large-scale proteome and organelle remodeling via selective autophagy. Mol Cell 81, 5082–5098 e5011

17. Cipolla, C. M., and Lodhi, I. J. (2017) Peroxisomal Dysfunction in Age-Related Diseases. Trends Endocrinol Metab 28, 297–308

18. Cho, D. H., Kim, Y. S., Jo, D. S., Choe, S. K., and Jo, E. K. (2018) Pexophagy: Molecular Mechanisms and Implications for Health and Diseases. Mol Cells 41, 55–64

19. Law, K. B., Bronte-Tinkew, D., Di Pietro, E., Snowden, A., Jones, R. O., Moser, A., Brumell, J. H., Braverman, N., and Kim, P. K. (2017) The peroxisomal AAA ATPase complex prevents pexophagy and development of peroxisome biogenesis disorders. Autophagy 13, 868–884

20. Soucy, T. A., Smith, P. G., Milhollen, M. A., Berger, A. J., Gavin, J. M., Adhikari, S., Brownell, J. E., Burke, K. E., Cardin, D. P., Critchley, S., Cullis, C. A., Doucette, A., Garnsey, J. J., Gaulin, J. L., Gershman, R. E., Lublinsky, A. R., McDonald, A., Mizutani, H., Narayanan, U., Olhava, E. J., Peluso, S., Rezaei, M., Sintchak, M. D., Talreja, T., Thomas, M. P., Traore, T., Vyskocil, S., Weatherhead, G. S., Yu, J., Zhang, J., Dick, L. R., Claiborne, C. F., Rolfe, M., Bolen, J. B., and Langston, S. P. (2009) An inhibitor of NEDD8-activating enzyme as a new approach to treat cancer. Nature 458, 732–736

21. Rusilowicz-Jones, E. V., Barone, F. G., Lopes, F. M., Stephen, E., Mortiboys, H., Urbe, S., and Clague, M. J. (2022) Benchmarking a highly selective USP30 inhibitor for enhancement of mitophagy and pexophagy. Life Sci Alliance 5

22. Marcassa, E., Kallinos, A., Jardine, J., Rusilowicz-Jones, E. V., Martinez, A., Kuehl, S., Islinger, M., Clague, M. J., and Urbe, S. (2018) Dual role of USP30 in controlling basal pexophagy and mitophagy. EMBO Rep 19, e45595

23. Riccio, V., Demers, N., Hua, R., Vissa, M., Cheng, D. T., Strilchuk, A. W., Wang, Y., McQuibban, G. A., and Kim, P. K. (2019) Deubiquitinating enzyme USP30 maintains basal peroxisome abundance by regulating pexophagy. J Cell Biol 218, 798–807

24. Katayama, H., Kogure, T., Mizushima, N., Yoshimori, T., and Miyawaki, A. (2011) A sensitive and quantitative technique for detecting autophagic events based on lysosomal delivery. Chem Biol 18, 1042–1052

25. Allen, G. F., Toth, R., James, J., and Ganley, I. G. (2013) Loss of iron triggers PINK1/ Parkin-independent mitophagy. EMBO Rep 14, 1127–1135

26. Walter, K. M., Schonenberger, M. J., Trotzmuller, M., Horn, M., Elsasser, H. P., Moser, A. B., Lucas, M. S., Schwarz, T., Gerber, P. A., Faust, P. L., Moch, H., Kofeler, H. C., Krek, W., and Kovacs, W. J. (2014) Hif-2alpha promotes degradation of mammalian peroxisomes by selective autophagy. Cell Metab 20, 882–897

27. Zhang, J., Tripathi, D. N., Jing, J., Alexander, A., Kim, J., Powell, R. T., Dere, R., Tait-Mulder, J., Lee, J. H., Paull, T. T., Pandita, R. K., Charaka, V. K., Pandita, T. K., Kastan, M. B., and Walker, C. L. (2015) ATM functions at the peroxisome to induce pexophagy in response to ROS. Nat Cell Biol 17, 1259–1269

28. Nazarko, T. Y., Ozeki, K., Till, A., Ramakrishnan, G., Lotfi, P., Yan, M., and Subramani, S. (2014) Peroxisomal Atg37 binds Atg30 or palmitoyl-CoA to regulate phagophore formation during pexophagy. J Cell Biol 204, 541–557

29. Bellot, G., Garcia-Medina, R., Gounon, P., Chiche, J., Roux, D., Pouyssegur, J., and Mazure, N. M. (2009) Hypoxia-induced autophagy is mediated through hypoxia-inducible factor induction of BNIP3 and BNIP3L via their BH3 domains. Mol Cell Biol 29, 2570–2581

30. Ordureau, A., Paulo, J. A., Zhang, J., An, H., Swatek, K. N., Cannon, J. R., Wan, Q., Komander, D., and Harper, J. W. (2020) Global Landscape and Dynamics of Parkin and USP30-Dependent Ubiquitylomes in iNeurons during Mitophagic Signaling. Mol Cell 77, 1124–1142 e1110

31. Wilhelm, L. P., Zapata-Munoz, J., Villarejo-Zori, B., Pellegrin, S., Freire, C. M., Toye, A. M., Boya, P., and Ganley, I. G. (2022) BNIP3L/NIX regulates both mitophagy and pexophagy. EMBO J, e111115

32. Stickle, N. H., Chung, J., Klco, J. M., Hill, R. P., Kaelin, W. G., Jr., and Ohh, M. (2004) pVHL modification by NEDD8 is required for fibronectin matrix assembly and suppression of tumor development. Mol Cell Biol 24, 3251–3261

33. Barghout, S. H., and Schimmer, A. D. (2021) E1 Enzymes as Therapeutic Targets in Cancer. Pharmacol Rev 73, 1–58

34. Hartwig, S., Knebel, B., Goeddeke, S., Koellmer, C., Jacob, S., Nitzgen, U., Passlack, W., Schiller, M., Dicken, H. D., Haas, J., Muller-Wieland, D., Lehr, S., and Kotzka, J. (2013) So close and yet so far: mitochondria and peroxisomes are one but with specific talents. Arch Physiol Biochem 119, 126–135

35. Andrade-Navarro, M. A., Sanchez-Pulido, L., and McBride, H. M. (2009) Mitochondrial vesicles: an ancient process providing new links to peroxisomes. Curr Opin Cell Biol 21, 560–567

36. Yamashita, S., Abe, K., Tatemichi, Y., and Fujiki, Y. (2014) The membrane peroxin PEX3 induces peroxisome-ubiquitination-linked pexophagy. Autophagy 10, 1549–1564

37. Bingol, B., Tea, J. S., Phu, L., Reichelt, M., Bakalarski, C. E., Song, Q., Foreman, O., Kirkpatrick, D. S., and Sheng, M. (2014) The mitochondrial deubiquitinase USP30 opposes parkin-mediated mitophagy. Nature 510, 370–375

38. Cunningham, C. N., Baughman, J. M., Phu, L., Tea, J. S., Yu, C., Coons, M., Kirkpatrick, D. S., Bingol, B., and Corn, J. E. (2015) USP30 and parkin homeostatically regulate atypical ubiquitin chains on mitochondria. Nat Cell Biol 17, 160–169

39. Liang, J. R., Martinez, A., Lane, J. D., Mayor, U., Clague, M. J., and Urbe, S. (2015) USP30 deubiquitylates mitochondrial Parkin substrates and restricts apoptotic cell death. EMBO Rep 16, 618–627

40. Gersch, M., Gladkova, C., Schubert, A. F., Michel, M. A., Maslen, S., and Komander, D. (2017) Mechanism and regulation of the Lys6-selective deubiquitinase USP30. Nat Struct Mol Biol 24, 920–930

41. Phu, L., Rose, C. M., Tea, J. S., Wall, C. E., Verschueren, E., Cheung, T. K., Kirkpatrick, D. S., and Bingol, B. (2020) Dynamic Regulation of Mitochondrial Import by the Ubiquitin System. Mol Cell 77, 1107–1123 e1110

42. Rusilowicz-Jones, E. V., Jardine, J., Kallinos, A., Pinto-Fernandez, A., Guenther, F., Giurrandino, M., Barone, F. G., McCarron, K., Burke, C. J., Murad, A., Martinez, A., Marcassa, E., Gersch, M., Buckmelter, A. J., Kayser-Bricker, K. J., Lamoliatte, F., Gajbhiye, A., Davis, S., Scott, H. C., Murphy, E., England, K., Mortiboys, H., Komander, D., Trost, M., Kessler, B. M., Ioannidis, S., Ahlijanian, M. K., Urbe, S., and Clague, M. J. (2020) USP30 sets a trigger threshold for PINK1-PARKIN amplification of mitochondrial ubiquitylation. Life Sci Alliance 3

43. Tsefou, E., Walker, A. S., Clark, E. H., Hicks, A. R., Luft, C., Takeda, K., Watanabe, T., Ramazio, B., Staddon, J. M., Briston, T., and Ketteler, R. (2021) Investigation of USP30 inhibition to enhance Parkin-mediated mitophagy: tools and approaches. Biochem J 478, 4099–4118

44. Zheng, J., Chen, X., Liu, Q., Zhong, G., and Zhuang, M. (2022) Ubiquitin ligase MARCH5 localizes to peroxisomes to regulate pexophagy. J Cell Biol 221

45. Clague, M. J., and Urbe, S. (2017) Integration of cellular ubiquitin and membrane traffic systems: focus on deubiquitylases. FEBS J 284, 1753–1766

46. Elcocks, H., Brazel, A. J., McCarron, K. R., Kaulich, M., Husnjak, K., Mortiboys, H., Clague, M. J., and Urbé, S. (2022) FBXL4 deficiency promotes mitophagy by elevating NIX. bioRxiv, 2022.2010.2011.511735

47. Montava-Garriga, L., Singh, F., Ball, G., and Ganley, I. G. (2020) Semi-automated quantitation of mitophagy in cells and tissues. Mech Ageing Dev 185, 111196

